# Developmental Expression of Transforming Growth Factor Induced Protein Promotes NF-Kappa-B Mediated Angiogenesis During Postnatal Lung Development

**DOI:** 10.1101/2020.05.28.121871

**Authors:** Min Liu, Cristiana Iosef, Shailaja Rao, Racquel Domingo-Gonzalez, Sha Fu, Paige Snider, Simon J. Conway, Gray S. Umbach, Sarah C. Heilshorn, Ruby E. Dewi, Mar J. Dahl, Donald M. Null, Kurt H. Albertine, Cristina M. Alvira

**Affiliations:** Stanford University, Department of Pediatrics, Center for Excellence in Pulmonary Biology; Liuyang People’s Hospital, Hunan, China; HB Wells Center for Pediatric Research, Indiana University School of Medicine, Indianapolis, IN 46202, USA; University of Texas Southwestern Medical School, University of Utah School of Medicine; Stanford University, Department of Materials Science & Engineering, University of Utah School of Medicine; Department of Pediatrics, University of Utah School of Medicine

**Author notes:** Address for Correspondence: Cristina M. Alvira, M.D., Stanford University School of Medicine, Center for Excellence in Pulmonary Biology, 770 Welch Road, Suite 435, Palo Alto, CA 94304. Contributions: ML, CI, SR, SF, GSU, RD, SCH, and CMA designed and executed studies, analyzed and interpreted results. PS and SJC created the TGFBI null mice. MJD, DMN, and KHA designed and executed the studies on preterm lambs. ML and CMA drafted the manuscript. All authors contributed edits and significant have given final approval for publication and agree to be accountable for the integrity of the information contained in this manuscript.

**Keywords:** Alveolarization, endothelial migration, colony stimulating factor-3, nitric oxide production, bronchopulmonary dysplasia

## Abstract

**Rationale:** Pulmonary angiogenesis is a key driver of alveolarization. Our prior studies showed that nuclear factor kappa-B (NFκB) promotes pulmonary angiogenesis during early alveolarization. However, the mechanisms regulating temporal-specific NFκB activation in the pulmonary vasculature are unknown.

**Objectives:** To identify mechanisms that activate pro-angiogenic NFκB signaling in the developing pulmonary vasculature.

**Methods:** Proteomic analysis of the lung secretome was performed using 2D-DIGE. NFκB activation and angiogenic function was assessed in primary pulmonary endothelial cells (PEC) and TGFBI-regulated genes identified using RNA-sequencing. Alveolarization and pulmonary angiogenesis was assessed in WT and TGFBI null mice exposed to normoxia or hyperoxia. Lung TGFBI expression was determined in premature lambs supported by invasive and noninvasive respiratory support.

**Measurements and Main Results:** Secreted factors from the early alveolar, but not the late alveolar or adult lung, promoted proliferation and migration in quiescent, adult PEC. Proteomic analysis identified transforming growth factor beta-induced protein (TGFBI) as a protein highly expressed by myofibroblasts in the early alveolar lung that promoted PEC migration by activating NFκB via αvβ3 integrins. RNA-sequencing identified *Csf3* as a TGFBI-regulated gene that enhances nitric oxide production in PEC. Loss of TGFBI in mice exaggerated the impaired pulmonary angiogenesis induced by chronic hyperoxia, and TGFBI expression was disrupted in premature lambs with impaired alveolarization.

**Conclusions:** Our studies identify TGFBI as a developmentally-regulated protein that promotes NFκB-mediated angiogenesis during early alveolarization by enhancing nitric oxide production. We speculate that dysregulation of TGFBI expression may contribute to diseases marked by impaired alveolar and vascular growth.

## Introduction

In contrast to many organs, significant lung development occurs postnatally. During alveolarization, the final stage of lung development, division of primitive airspaces by secondary septation, and exponential growth of the pulmonary microvasculature by angiogenesis markedly increases gas exchange surface area (1). As the lung matures, alveolarization and angiogenesis slows, and the pulmonary endothelium transitions from an activated to a quiescent phenotype characteristic of the mature vasculature (2).

Pulmonary angiogenesis is a key driver of alveolarization. Inhibiting angiogenesis impairs alveolarization, while enhancing angiogenesis preserves alveolarization during injury (3-5). Dysregulated angiogenesis is observed in premature infants with bronchopulmonary dysplasia (BPD), a chronic lung disease characterized by impaired alveolarization that represents the most common complication of extreme prematurity (6). The extension of alveolarization into postnatal life provides an important window of opportunity for lung repair and regeneration (7). Thus, elucidating pathways that promote pulmonary angiogenesis and alveolarization have important clinical implications.

We previously showed that endogenous NFκB activation promotes pulmonary angiogenesis in the early alveolar lung (8). Blocking NFκB in early alveolar pulmonary endothelial cells (PEC) impairs angiogenic function, and pharmacologic inhibition of NFκB in young mice impairs alveolar and vascular growth yet has no effect on adult mice. However, the mechanisms that induce temporal-specific activation of NFκB in the developing pulmonary vasculature remain unknown.

The tissue microenvironment modulates angiogenesis in development and disease. During early lung development, the alveolar epithelium regulates vascular patterning by expressing VEGF to promote EC survival, proliferation, and migration. In cancer, activated stromal fibroblasts develop a myofibroblast phenotype, co-localize with tumor vasculature, and promotes angiogenesis by expressing growth factors and modulating the extracellular matrix (9). Although myofibroblasts are required for alveolarization (10), whether they function to regulate pulmonary angiogenesis has not been explored.

In this study, we hypothesized that unique factors in the early alveolar microenvironment induce temporal-specific activation of NFκB in the pulmonary endothelium. We profiled the lung secretome during development and identified transforming growth factor beta-induced protein (TGFBI) as protein highly expressed by myofibroblasts during early alveolarization. We show that TGFBI induces NFκB activation through αvβ3 integrins, increasing nitric oxide (NO) production by increasing the expression of colony stimulating factor-3 (*Csf3*), an NFκB downstream target gene. TGFBI null mice exhibit decreased pulmonary vascular density, and a marked impairment of alveolar and vascular growth in repose to chronic hyperoxia. Further, TGFBI expression was dysregulated in a premature lamb model of disrupted alveolarization. Together, our data identify a novel myofibroblast-endothelial cell axis that serves to guide pulmonary angiogenesis during early alveolarization and implicate a role for dysregulated TGFBI in the pathogenesis of BPD.

## Methods

Please see the Online Methods for full details.

### Animal Models

C57BL/6 neonatal mice at early alveolarization (P6) and adult mice were purchased from Charles River Lab. TGFBI-/-mice have been described previously (11). Mice containing an endothelial cell specific deletion of IKKβ were generated by crossing IKKβ^fl/fl^ mice (12) with *Pdgfb*-iCre mice (13). For hyperoxia experiments, litters of P0 pups were maintained in room air (normoxia) or 80% O_2_ (hyperoxia) for 14 days (14). Lung morphometric analysis was performed as previously described (8, 15).

The methods for delivery and management of chronically ventilating preterm lambs are reported (16-19). Time-pregnant ewes at 132 ± 2 d of gestation (term ∼150 d gestation) were used. At ∼3h of age, the preterm lambs were randomized to IMV or NRS, as previously described (18) for a total of 21d. Control lambs were born at term.

Protocols for the animal studies adhered to American Physiological Society/US National Institutes of Health guidelines for humane use of animals for research and were prospectively approved by the Institutional Animal Care and Use Committee at Stanford University and the University of Utah Health Sciences Center.

### Lung Conditioned Media

Lung conditional medium (LCM) was prepared from lung tissue from C5BL/6 mice at the early alveolar (P6), late alveolar (P16) and adult (8-10 weeks) stages of development (20), and proteins analyzed by 2D DIGE protein expression profiling.

### Western Immunoblot and Immunofluorescence

Whole cell protein lysates were extracted from lung tissue and western blot performed (15). Immunostaining was performed on formalin-fixed or frozen lung sections (8), probed with primary antibodies against CD31, TGFBI, NFκB p65 or von Willebrand factor.

### Isolation of Primary Pulmonary Endothelial Cells (PEC)

PEC were isolated from P6 or adult C57BL/6, *Tgfbi*^(-/-)^and *Tgfbi* ^(+/+),^ and *Pdgfb*-iCre^(+/-)^ IKKβ^fl/fl^ mice as described previously (8, 15). Cells from passage 0-2 were used for all assays as described in the Supplemental Methods. TGFBI neutralization was performed with anti-TGFBI antibodies (4 μg/ml), TGFBI stimulation with recombinant TGFBI (10 μg/ml), NFκB inhibition with the pharmacologic inhibitor, BAY11-0782 (2.5 μM), and αvβ3 inhibition with signaling anti-αvβ3 integrin (4 μg/ml) antibodies.

### RNA interference

PEC were transfected NTC, integrin αV, integrin β3, or Csf3 On-Target Plus SMART pool siRNA using Lipofectamine 2000 for 6h as previously described (21).

### RNA-Seq Analysis

Total RNA was extracted and RNA-sequencing performed by Quick Biology. Genes showing altered expression with P< 0.05 and more than 1.5-fold change were considered differentially expressed.

### Measurement of NO in PEC

NO production was determined by loading the cells with 4-amino-5-methylamino-2’,7’-difluorofluorescein diacetate (DAF-FM) (22) prior to immunofluorescent imaging as previously described (23).

### Statistics

Statistical differences between two groups were determined by Student’s t-test, One- or Two-Way ANOVA as appropriate. A P value of ≤0.05 was considered statistically significant.

## Results

### Factors secreted by the early alveolar lung activate pro-angiogenic pathways in adult PEC

To determine if factors present in the early alveolar microenvironment activate NFκB and modulate PEC angiogenic function, we collected lung condition medium (LCM) from mice at different stages of development and assessed NFκB activation and PEC migration. Under control conditions, NFκB subunits are constitutively expressed, but only translocate to the nucleus upon activation. At baseline, adult PEC demonstrated minimal active NFκB (Fig. 1A). However, incubation with early alveolar LCM increased NFκB activation by 2.65-fold (P<0.001) (Fig. 1A and B). In contrast, incubation with late alveolar LCM increased NFκB activation only slightly (P<0.05), and adult LCM had no effect. Similarly, adult PEC migrated slowly when cultured in starvation media (Fig. 1C). The early alveolar LCM was as effective as 5% FBS in inducing adult PEC migration, resulting in 43% scratch closure (P<0.0001). In contrast, the adult LCM induced migration only minimally (P<0.05). Taken together, these data suggested that factors present in the early alveolar lung microenvironment can induce NFκB activation and promote migration in adult PEC.

**Figure 1:**
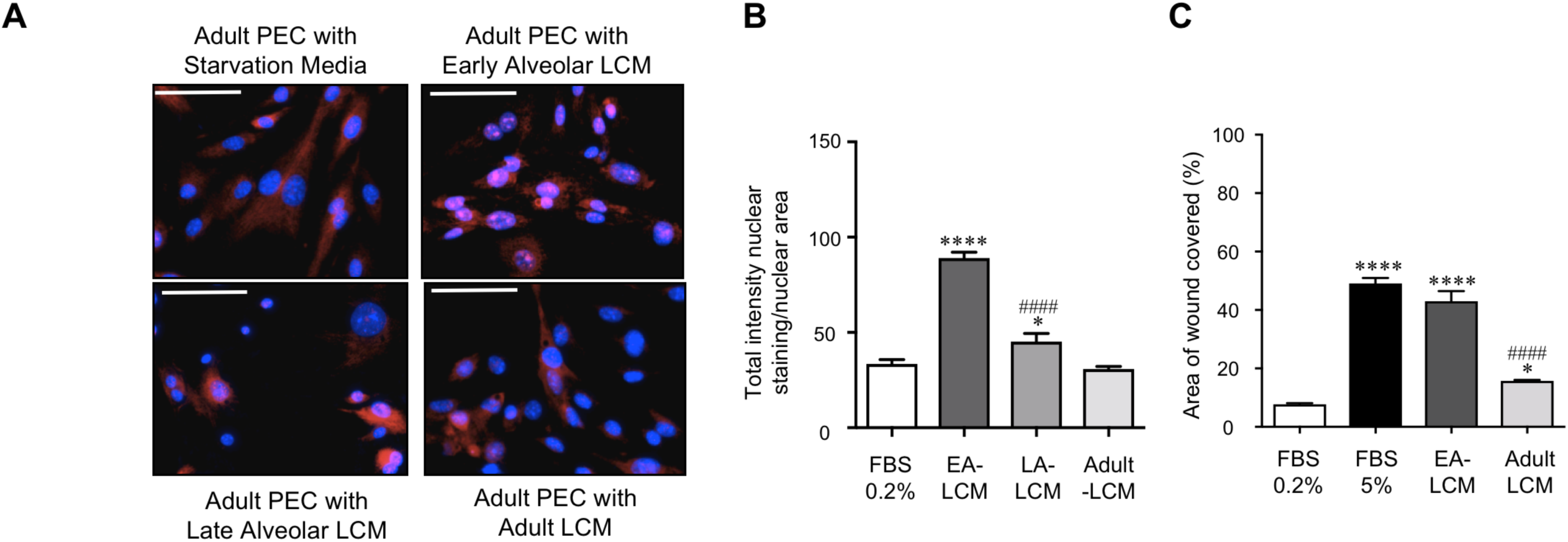
Factors secreted by the early alveolar lung activate pro-angiogenic pathways in adult pulmonary endothelial cells. (A) Representative immunofluorescent images of adult PEC incubated with starvation media, or early alveolar (EA), late alveolar (LA) or adult lung conditioned media (LCM) for 24h followed by immunostaining to detect the NFκB subunit, p65 (red) and chromatin (blue). (B) Quantification of the total intensity of nuclear p65 with *P<0.05, and ****P<0.0001 vs. starvation, and ####P<0.0001 vs. EA-LCM, with n=6 per group. (C) EC scratch assays performed using adult PEC incubated with starvation media, 5% FBS, EA-LCM, and adult LCM, and the percent scratch area covered at 24h calculated. ****P<0.0001 vs. starvation, and ####P<0.0001 vs. EA-LCM, with n=3 per group.

### TGFBI is highly expressed in the early alveolar lung but absent in the adult lung

To identify factors uniquely present in the early alveolar lung microenvironment, we compared all of the secreted proteins in the three LCM by two-dimensional difference gel electrophoresis (2-D DIGE) (Fig. E1), and identified 20 proteins that were highly expressed in the early alveolar lung secretome by mass spectrometry (Table E1). Of this group we selected transforming growth factor induced protein (TGFBI) for further investigation, a classically secreted protein that has recently been shown to be highly expressed by myofibroblasts by single cell RNA sequencing of the developing mouse lung (24). We confirmed higher expression of TGFBI in the early alveolar LCM (P<0.001, Fig.2 A), and in agreement with previous data (11), highest TGFBI protein in whole lung during early alveolarization, followed by an age-dependent decrease over time (Fig. 2B). Immunostaining of early alveolar lung tissue identified numerous cells with intense TGFBI expression (Fig. 2C), located at the tips of secondary septa, characteristic locations for alveolar myofibroblasts. In contrast, TGFBI immunoreactivity was completely absent in the adult lung. Similar findings were observed in the lungs of lambs (Fig. 2D), with high TGFBI expression in cells at the septal tips at 1 day of life, corresponding to early alveolarization, but reduced TGFBI expression by three weeks of age.

**Figure 2:**
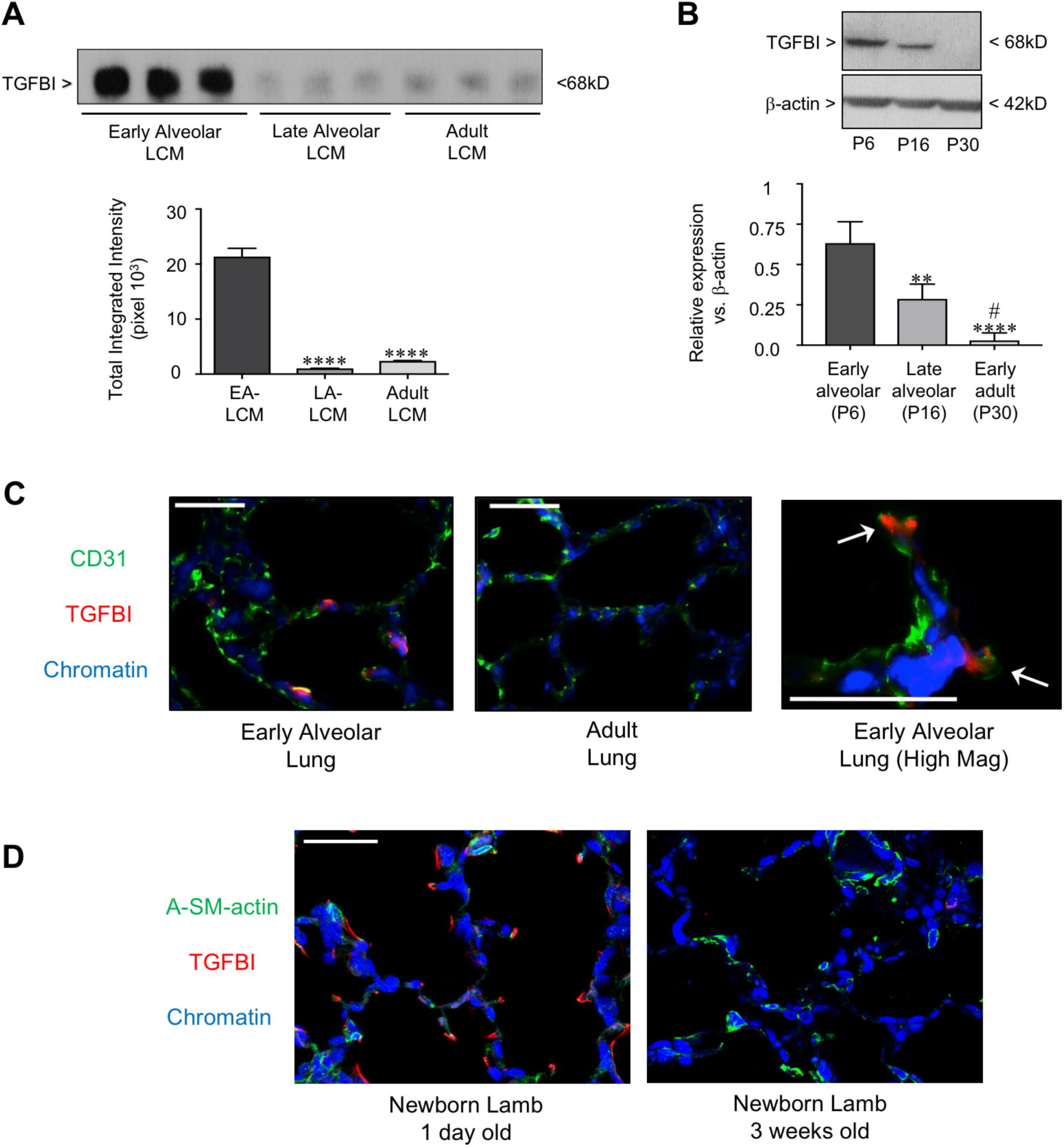
TGFBI is highly expressed in the early alveolar lung but absent in the adult lung. (A) Western blot to determine TGFBI protein in the EA-, LA-, and adult LCM. ****P<0.0001 vs. EA=LCM, with n=3. (B) Western blot to determine TGFBI protein relative to β-actin in whole lung from mice at the early alveolar (P6), late alveolar (P16), and adult (P30) stages of development. **P<0.01 and ****P<0.0001 vs. P6, and #P<0.05 vs. P16, with n=4 per group. (C) Representative images obtained from lung cryosections obtained from P6 and adult mice to detect CD31 (green), TGFBI (red) and chromatin (blue). Arrows point to TGFBI-positive cells at tips of secondary septa. Calibration mark=100μm. (D) Representative images obtained from lung tissue from lambs at the early alveolar (Day 1) and late alveolar (3 week) stages of development to detect α-smooth muscle-actin (green), TGFBI (red) and chromatin (blue). Calibration mark=50μm.

### TGFBI is necessary and sufficient for promoting early alveolar and adult PEC migration

We next determined whether TGFBI was required for the early alveolar LCM to enhance adult PEC migration. The addition of anti-TGFBI antibodies, but not isotype control IgG, significantly impaired the capacity of the early alveolar LCM to promote adult PEC migration (P<0.0001, Fig.3A), but had no effect on its proliferative effect (Fig. 3B). To determine whether TGFBI was sufficient to promote PEC migration, we employed recombinant TGFBI (rTGFBI) in microfluidic chemotaxis assays that permitted the creation of stable, linear gradients of chemotactic agents. Early alveolar PEC exposed to a gradient of starvation media migrated randomly, while those exposed to a gradient of VEGF demonstrated directed migration toward the source (P<0.0001) (Fig. 3C). Both the early alveolar LCM and rTGFBI induced similar directed migration. Moreover, rTGFBI promoted migration of both the early alveolar and adult PEC in endothelial scratch assays and Boyden chamber assays (Fig. E2).

**Figure 3:**
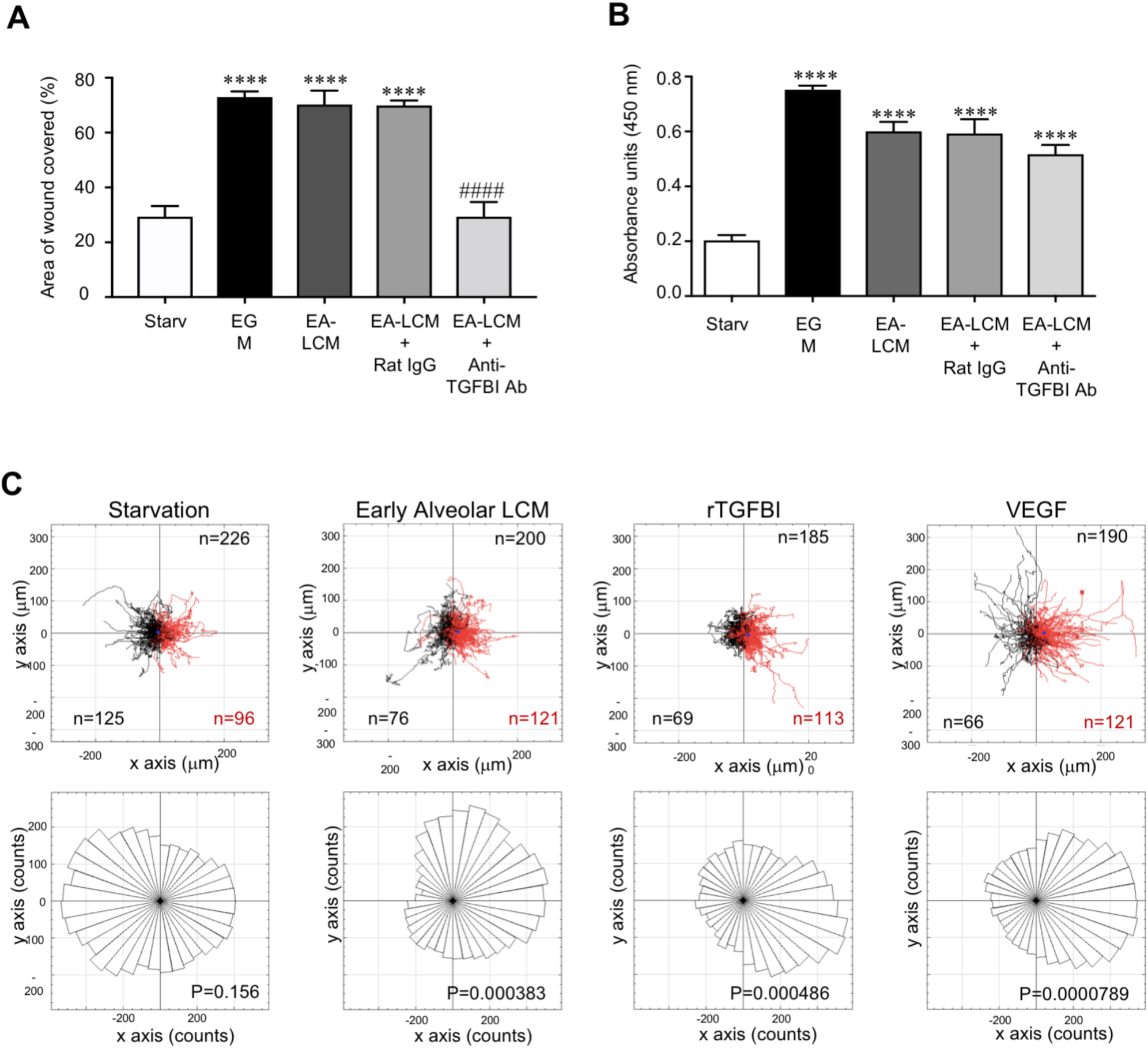
TGFBI is necessary and sufficient to promote PEC migration. (A) Endothelial scratch assays were performed using adult PEC incubated with starvation media, 5% FBS, EA-LCM, EA-LCM plus isotype control IgG, and EA-LCM plus anti-TGFBI antibodies (4 μg/ml) and the percent scratch area covered at 24h calculated. ****P<0.0001 vs. starvation, and ####P<0.0001 vs. EA-LCM, with n=3 per group. Representative results from three separate experiments. (B) BrdU incorporation assays to assess adult PEC proliferation at 24h in cells incubated with 5% FBS, EA-LCM, EA-LCM + IgG, and EA-LCM + anti-TGFBI antibodies. ****P<0.0001 vs. starvation with n=3-6 per group. (C) Tracks of individual cells (top) and directional histograms (bottom) from live cell imaging and tracking of early alveolar PEC subjected to microfluidic chemotaxis assays performed with starvation media, EA-LCM, starvation +TGFBI or starvation + VEGF (50 ng/ml) with each chamber containing a source on the right side and a sink on the left. Total number of individual cells tracked is reported in the upper left corner, and the number of cells migrating away (black) or toward (red) in the bottom left and right corners respectively. In each group there were between 3-5 cells with a migration of net zero, accounting for the remaining cells making up the total n number. P value is shown on the image.

### TGFBI-mediated PEC migration is NFκB-dependent

We next assessed whether the pro-migratory effect of TGFBI is NFκB-dependent. Similar to the migration results, early alveolar LCM containing control IgG increased NFκB activation by 80% (P<0.001), but the addition of anti-TGFBI antibodies significantly blunted this effect (P<0.05) (Fig. 4A). rTGFBI significantly increased NFκB activity in early alveolar PEC (Fig. 4B, P<0.001), and increased NFκB-DNA binding as early as 30 min, with peak NFκB-DNA binding observed at 1h (Fig. 4C). In addition, inhibiting NFκB with BAY-7082 (25), completely abrogated TGBFI-mediated migration (P<0.0001, Fig. 4D). Further, we performed studies using PEC obtained from mice containing an endothelial-specific deletion of IKKβ, the primary activator of NFκB in early alveolar PEC (21). Although rTGFBI increased migration in WT PEC (P<0.01), TGFBI-induced migration was absent in PEC lacking IKKβ. Taken together, these data demonstrate that TGFBI-mediated migration is IKKβ/NFκB dependent (Fig. 4E).

**Figure 4:**
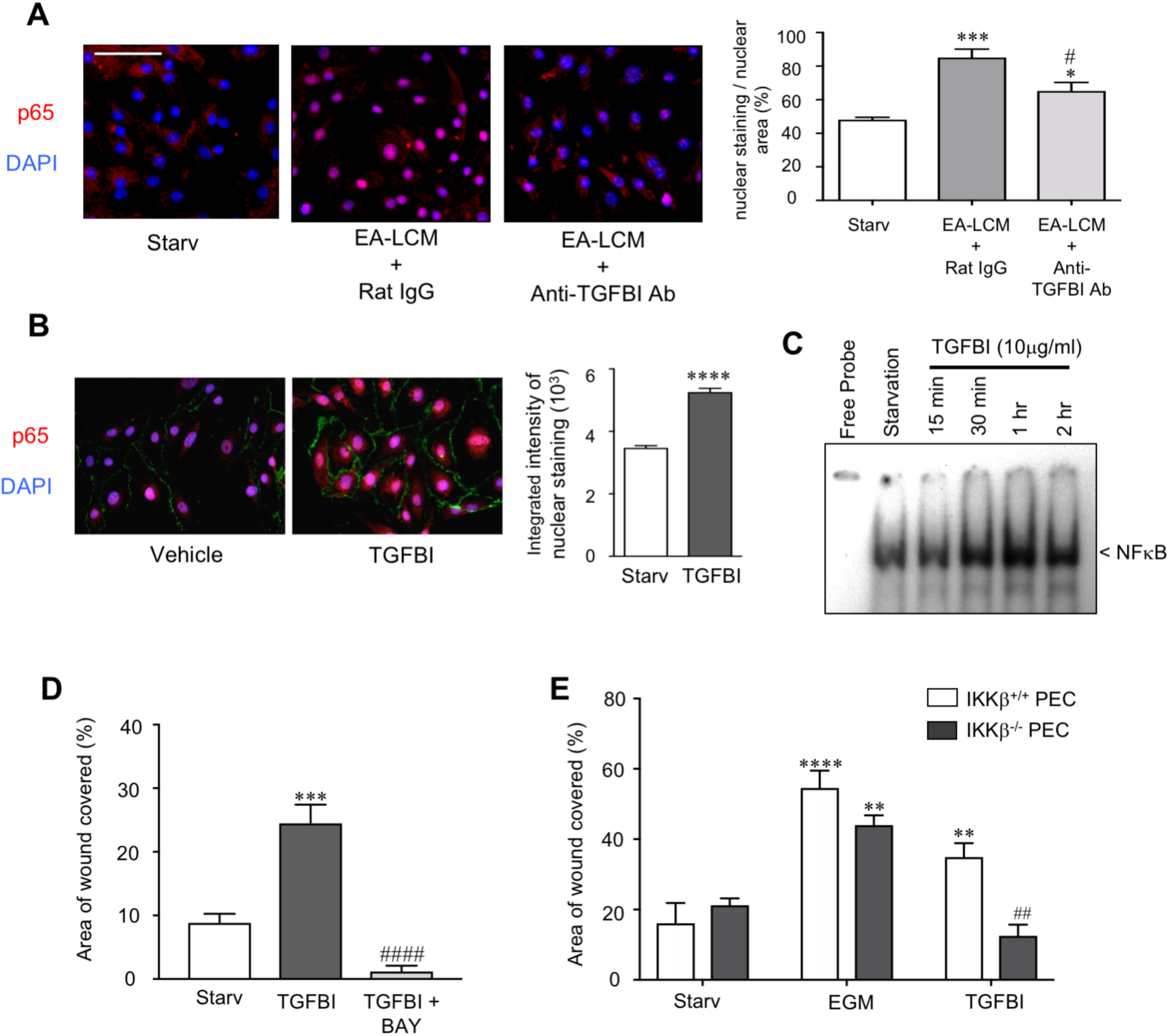
TGFBI-mediated pulmonary endothelial cell migration is NFκB-dependent. (A) Representative images of adult PEC incubated with starvation media, EA-LCM + IgG, and EA-LCM + anti-TGFBI antibodies for 24h followed by immunostaining to detect p65 (red) and chromatin (blue), with quantification to determine the total intensity of nuclear p65 over nuclear area. *P<0.05, and ***P<0.001 vs. starvation, and #P<0.05 vs. EA-LCM + IgG, with n=5 per group. (B) Representative images from early alveolar PEC incubated with starvation media + vehicle or starvation media + rTGFBI for 24h stained to detect the p65 (red), CD31(green), and DAPI (blue), with quantification of total intensity of nuclear p65. ****P<0.0001 with n=127 control cells and n=112 rTGFBI-stimulated cells. (C) Representative EMSA to detect NFκB-DNA binding in early alveolar PEC exposed to starvation media, or starvation media + TGFBI. (D) Scratch assays using early alveolar PEC stimulated with starvation media, starvation media + rTGFBI, or starvation media +rTGFBI and BAY 11-7082 (2.5 μM), with the percent scratch area covered at 24h calculated. ***P<0.001 vs. starvation and ####P<0.0001 vs. rTGFBI, with n=4 per group. (E) Scratch assays performed using wild type PEC (IKKβ^+/+^) and PEC lacking the NFκB activator, IKKβ (IKKβ^-/-^) stimulated with starvation media, EGM, starvation media + rTGFBI and the percent scratch area covered at 24h calculated. **P<0.01 and ****P<0.0001 vs. starvation and #P= 0.003 vs. TGFBI IKKβ^+/+^, with n=4 per group.

### TGFBI–mediated NFκB activation and endothelial migration requires avβ3 integrins

We next performed studies to identify how TGFBI was mediating these effects. TGFBI contains a carboxy-terminal Arg-Gly-Asp (RGD) sequence that allows binding to integrins (26). We focused initially on αvβ3, an integrin up-regulated in angiogenic vascular tissue (27), that activates NFκB in EC (28, 29). rTGFBI significantly increased NFκB activity in early alveolar PEC pre-treated with control IgG but had no effect on on PEC treated with anti-αvβ3 integrin antibodies (Fig. 5A and B). Although both rTGFBI alone and rTGFBI+IgG promoted PEC migration to a similar degree (Fig. 5C), anti-αvβ3 antibodies completely blocked rTGFBI-induced PEC migration. Finally, we transfected early alveolar PEC with NTC, αv integrin, or β3 integrin siRNA. In vehicle-stimulated cells, migration was similar between the three groups (Fig. 5D). rTGFBI significantly increased migration in the NTC-transfected PEC (P<0.01) but did not increase migration in PEC transfected with either αv or β3 siRNA.

**Figure 5:**
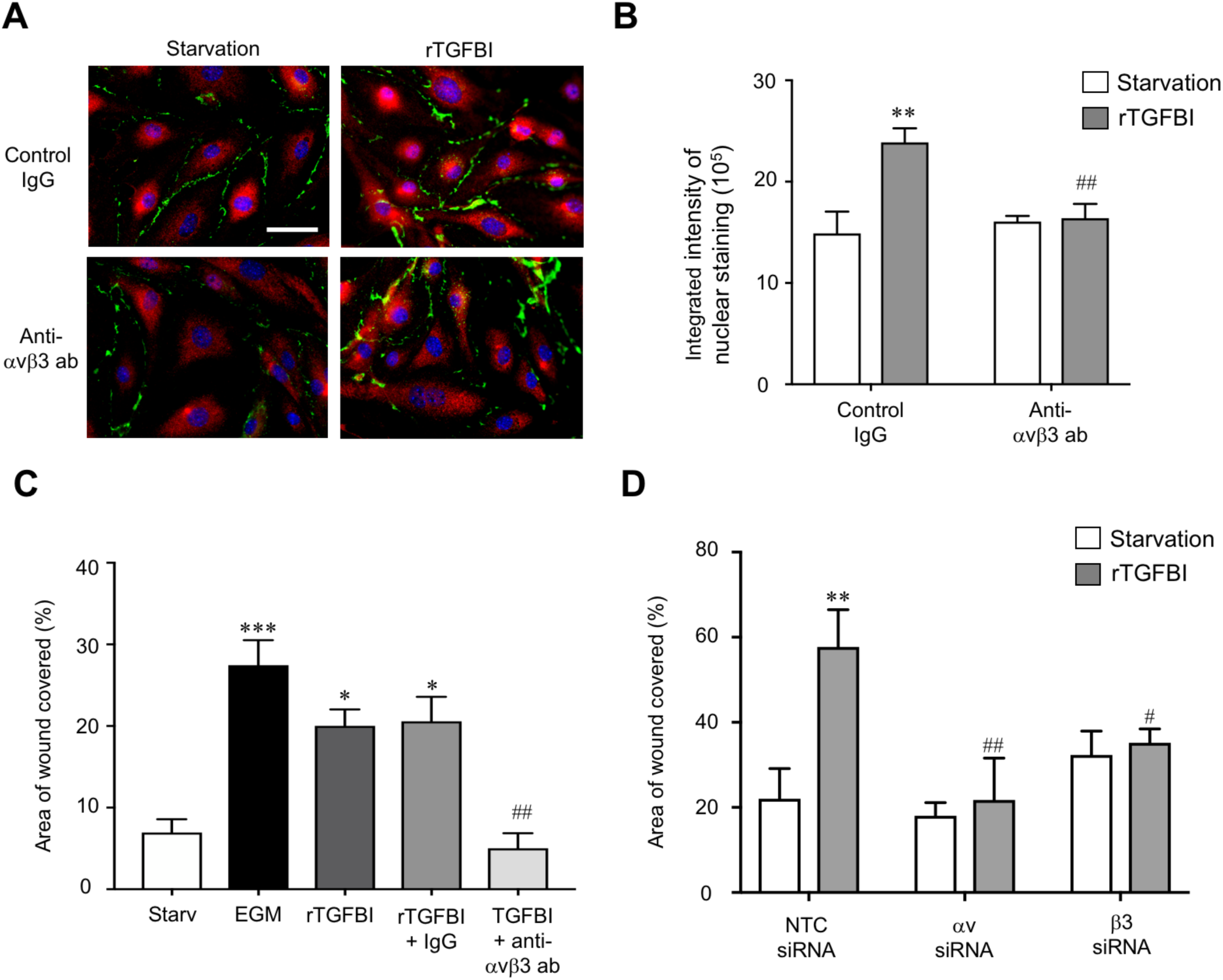
TGFBI–mediated endothelial migration requires αvβ3 integrins. (A) Representative images obtained from early alveolar PEC pretreated with either IgG or anti-αvβ3 antibodies prior to stimulation with starvation media + vehicle or starvation media + rTGFBI for 24h stained to detect the p65 (red), CD31(green), and DAPI (blue), with (B) quantification of total intensity of nuclear p65. **P<0.01 versus starvation + IgG, and ## versus rTGFBI + IgG with n=4. (C) Scratch assays with early alveolar PEC incubated with starvation media, EGM, starvation media + rTGFBI, starvation media + rTGFBI + IgG, and starvation media + rTGFBI plus anti-αvβ3 antibodies and the percent scratch area covered at 24h calculated. *P<0.05 and ***P<0.001 vs. starvation and ##P<0.01 vs. starvation + rTGFBI+ IgG, with n=3 per group. Representative result from 2 independent experiments. (D) Scratch assays were performed using early alveolar PEC transfected with NTC, integrin αv, and integrin β3 siRNA. At 48h post transfection, the groups were incubated with starvation media, EGM, starvation media + rTGFBI and the percent scratch area covered at 24h calculated. **P<0.01 vs. starvation, and #P<0.05 and ##P<0.01 vs. NTC stimulated with rTGFBI, with n=3-4 per group. Representative result from 4 independent experiments.

### TGFBI increases Csf3, a modulator of NO, in early alveolar PEC

To identify mechanism by which TGFBI promotes PEC migration, we profiled TGFBI-responsive genes using RNA-Seq. Given that rTGFBI stimulated both early alveolar and adult PEC migration (Fig. E2), we looked for genes induced by rTGFBI in both groups. Hierarchical clustering of differentially expressed genes demonstrated good clustering of vehicle- and rTGFBI-stimulated samples (Fig. 6A). rTGFBI significantly dysregulated 56 genes in early alveolar (Table E2) and 64 genes in adult PEC (Table E3), however, only 3 genes were shared (Fig. 6B). Colony stimulating factor-3 (*Csf3)*, a known NFκB- target gene (30), was among the shared genes, upregulated 3.01-fold in early alveolar and 2.77-fold in adult PEC by rTGFBI (Fig. 6C). We confirmed that rTGFBI induced a 3.4-fold increase in *Csf3* gene expression in early alveolar PEC by qPCR (Fig. 6D), and increased CSF3 protein (Fig. 6E). Prior studies found that CSF3 promotes EC migration by increasing in NO (31). Therefore, we loaded cells with the NO sensitive dye, DAF-FM diacetate prior to stimulation with vehicle or rTGFBI and found that rTGFBI increased NO in the PEC by more than 2-fold compared to vehicle (Fig. 6F).

**Figure 6:**
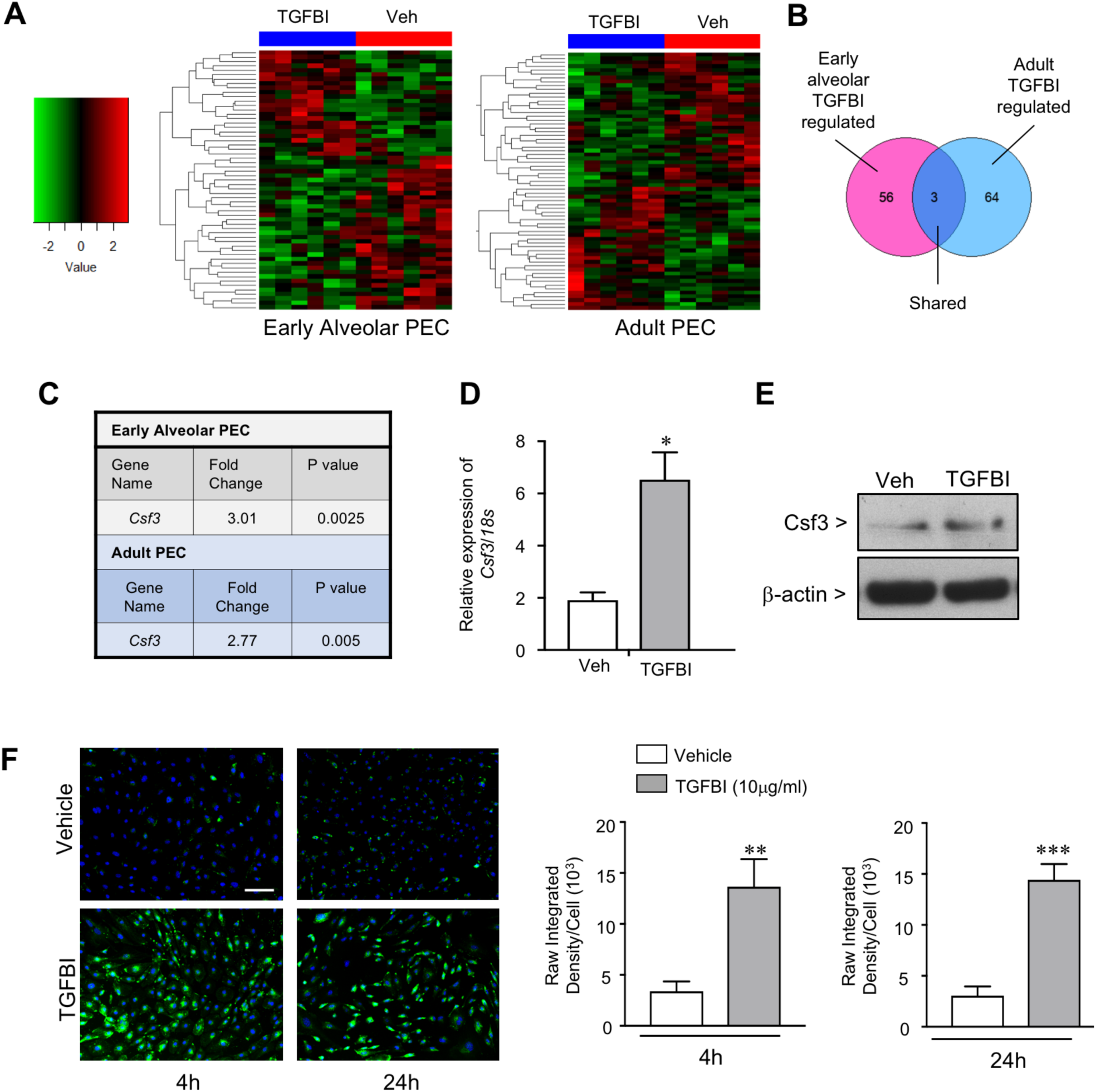
TGFBI increases the expression of Csf3, a modulator of nitric oxide, in early alveolar PEC. (A) Heat map of differentially expressed genes identified by RNA-Seq in early alveolar and adult PEC stimulated with vehicle or rTGFBI for 6h. Up-regulated genes are in red and down-regulated genes in green. (B) Venn diagram of unique and shared TGFBI-regulated genes in early alveolar and adult PEC. (C) *Csf3* was a gene up-regulated by rTGFBI in both early alveolar and adult PEC. (D) Gene expression of *Csf3* was determined by qRT-PCR in early alveolar PEC stimulated with vehicle or rTGFBI for 6h. *P<0.05 vs. vehicle with n=4 per group. (E) Representative western blot to detect CSF3 protein expression in whole cell lysates obtained from early alveolar PEC stimulated with vehicle or rTGFBI for 4hr. (F) NO production assays in early alveolar PEC stimulated with starvation media containing vehicle or rTGFBI for 4hr and 24h, and loaded with the NO sensitive dye, DAF-FM. Representative images taken to detect NO (green) and chromatin (blue). Calibration mark=100µm. Quantification of the raw integrated density of NO fluorescent signal per cell in early alveolar PEC stimulated with vehicle or rTGFBI for 4h and 24 hr, respectively, with **P<0.01, and ***P<0.0001 vs. vehicle, with n=5-8 per group. Results are representative of 3 independent experiments.

### Silencing Csf3 abrogates the TGFBI-mediated induction of NO and migration of early alveolar PEC

To determine if the TGFBI-mediated effects require *Csf3*, we performed additional studies where we silenced *Csf3*. Transfection of early alveolar PEC with *Csf3* siRNA effectively reduced CSF3 protein expression by 42h (Fig. 7A). rTGFBI increased migration 2-fold in NTC siRNA-transfected cells (Fig. 7B) but did not significantly enhance migration in *Csf3* siRNA-transfected cells. rTGFBI also enhanced NO production in NTC siRNA-transfected cells (Fig. 7C). However, rTGFBI-mediated increases in NO were completely blocked with *Csf3* silencing. Taken together, these results demonstrate that TGFBI promotes PEC migration by augmenting *Csf3*-dependent NO production.

**Figure 7:**
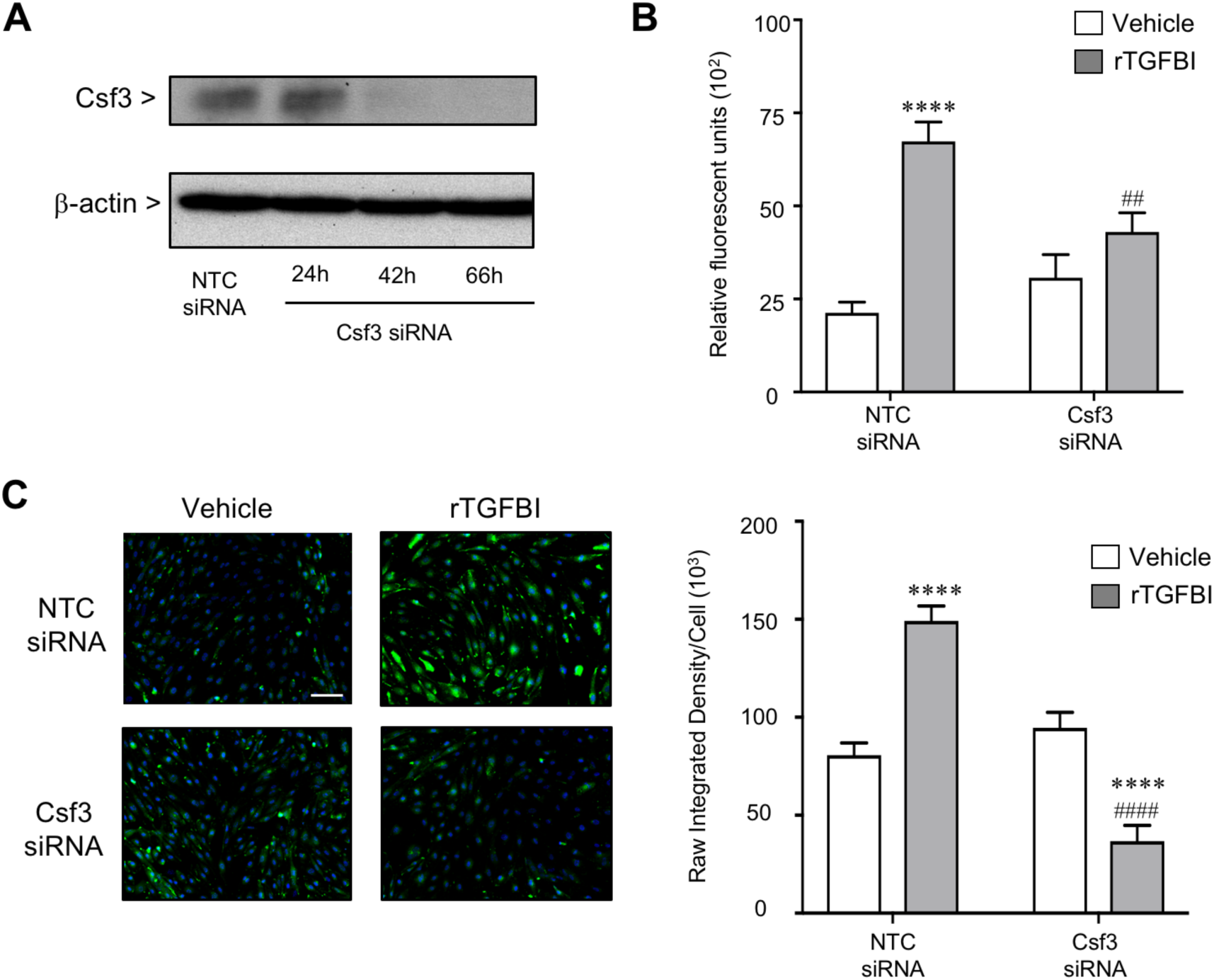
Silencing Csf3 abrogates the TGFBI-mediated induction of NO and migration of early alveolar PEC. (A) Representative western blot to detect Csf3 protein expression relative to β-actin in early alveolar PEC transfected with NTC and Csf3 siRNA. (B) Boyden chamber assays to detect chemotactic migration in early alveolar PEC transfected with NTC or Csf3 siRNA incubated at 48h after transfection with starvation media or starvation media containing rTGFBI for 8h. ****P <0.0001 vs. vehicle NTC siRNA and ##P<0.01 vs. NTC siRNA TGFBI, with n=10-12 replicates. Results are a representative example of three independent experiments. (C) NO production assays in early alveolar PEC transfected with NTC or Csf3 siRNA and stimulated with starvation + rTGFBI for 24h. Representative images to detect NO (green) and chromatin (blue). Quantification of the raw integrated density of NO fluorescence per cell in early alveolar PEC stimulated with vehicle or rTGFBI for 24h. ****P<0.0001 vs. vehicle, and #^###^P<0.0001 vs. rTGFBI stimulated NTC transfected.

### Loss of TGFBI impairs pulmonary vascular growth in mice, and TGFBI expression is dysregulated in preterm lambs with impaired alveolarization

To assess the physiological role of TGFBI in alveolarization and pulmonary angiogenesis, we evaluated mice containing a global deletion of TGFBI in normoxia and in response to chronic hyperoxia, a stimulus that disrupts pulmonary angiogenesis and alveolarization (32). These mice were reported to have impaired alveolarization at baseline, but abnormalities in vascular growth were not observed. In keeping with prior results, TGFBI null mice (TGFBI^-/-^) exhibited a 20% decrease in radial alveolar count (P<0.0001) and a 128% increase in distal airspace area (P<0.0001) compared to WT mice (Fig. 8A-C). As expected, chronic hyperoxia disrupted alveolarization in the WT mice, but induced a more exaggerated phenotype in the TGFBI^-/-^ mice, reducing radial alveolar count by almost 70% (P<0.0001) and further increased the already dilated distal airspaces (P<0.001). Under control conditions, TGFBI null mice exhibited a 33% reduction in pulmonary vascular density as compared to WT (P<0.0001) (Fig. 8D and E). Chronic hyperoxia reduced pulmonary vascular density in WT mice by 46% (P<0.0001), but caused a more exaggerated disruption of pulmonary vascular growth in TGFBI^-/-^ mice, decreasing pulmonary vascular density by 70% (P<0.0001), resulting in an almost 3-fold reduction in distal vessels in TGFBI^-/-^ compared to WT mice. Taken together, these results demonstrate that TGFBI is required for physiologic pulmonary vascular growth, and that loss of TGFBI worsens the detrimental effects of chronic hyperoxia on alveolarization and angiogenesis.

**Figure 8.**
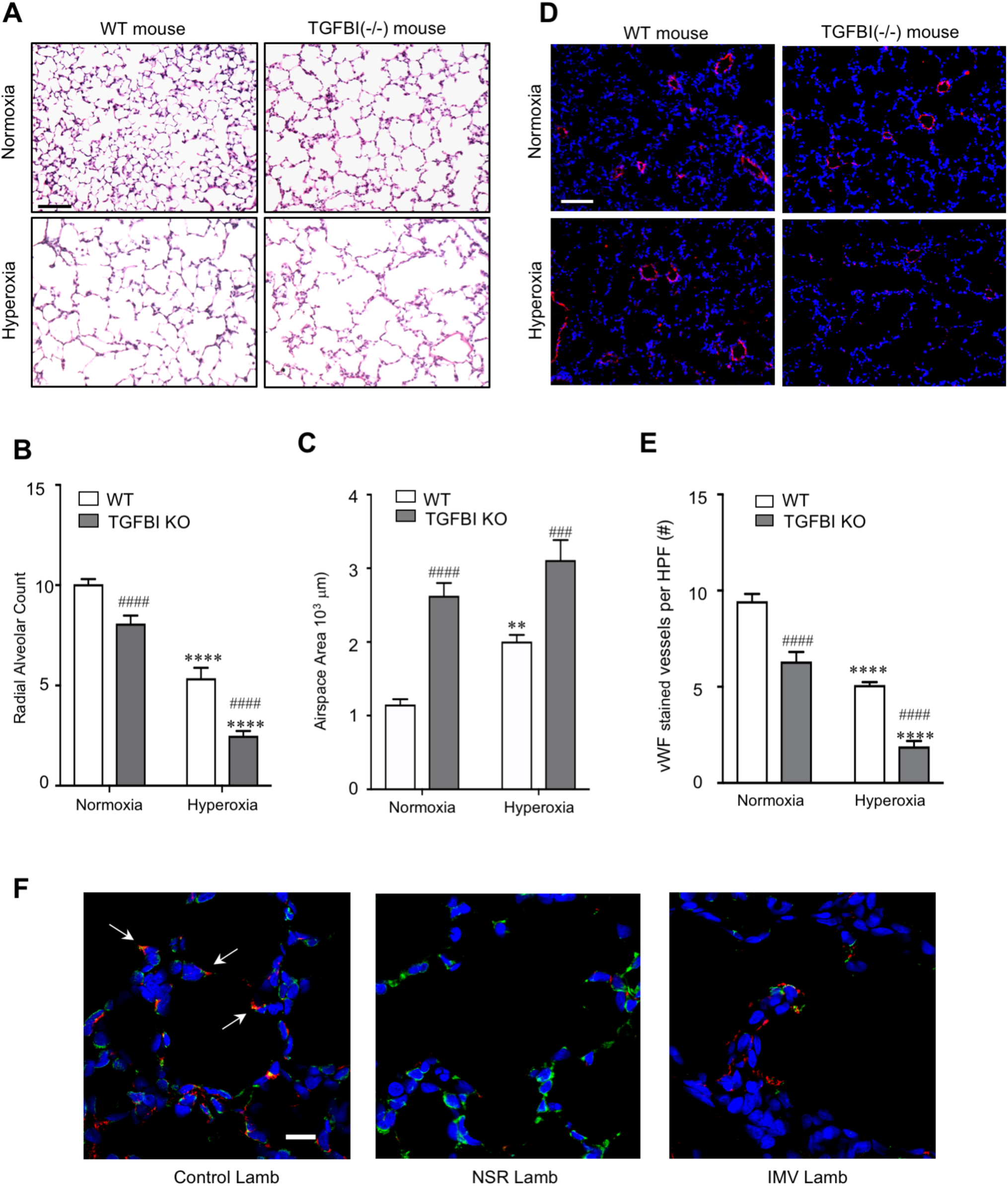
Loss of TGFBI in mice impairs pulmonary parenchymal and vascular growth and TGFBI expression is dysregulated in experimental models of impaired alveolarization. (A) Representative images obtained from P14 wild type (WT) and TGFBI^-/-^ mice maintained in normoxia or chronic hyperoxia (80% O_2_ from P1-P14). Calibration bar=100µm. Quantification of radial alveolar counts (B) and (C) distal airspace area. **P<0.01 and ****P<0.0001 vs. normoxia, and ###P<0.001 and ####P<0.0001 vs. WT via 2-WAY ANOVA, with n=4-7 per group. (D) Representative images stained to detect vWF (red) and DAPI (blue). Calibration bar=100µm. (E) Quantification of vWF-stained vessels less than 100μm per high powered field in 20 non-overlapping sections per mouse. ****P>0.0001 vs. normoxia, and ####P<0.0001 vs. WT with n=5-7 mice per group. (F) Representative images obtained from control term newborn lambs, and premature lambs treated with noninvasive respiratory support (NRS) or invasive mechanical ventilation (IMV), stained to detect TGFBI (red), alpha-smooth muscle actin (green) and chromatin (blue). Calibration bar=20 µm.

Finally, we explored whether TGFBI expression was altered in a large animal model of BPD, where preterm lambs are supported with either non-invasive respiratory support (NRS) or invasive mechanical ventilation (IMV). Control, term lambs had focal staining of TGFBI throughout the lung, including high expression at the tips of all the secondary septa (Fig. 8F, arrows). Preterm lambs supported with NRS exhibited many, thin secondary septa, and a marked reduction in TGFBI immunostaining. Preterm lambs supported by IMV, however, had abnormally thickened secondary septa, with a heightened expression but abnormal localization of TGFBI along the length of the thick secondary septa rather than the normal localization at the septal tips.

## Discussion

During early postnatal life, growth of the pulmonary vasculature serves as a driver of alveolarization. In this study, we explored the mechanisms that activate pro-angiogenic NFκB signaling in the pulmonary endothelium during early alveolarization. We identified TGFBI, as a secreted protein highly expressed in early alveolarization, corresponding to the time when NFκB is endogenously active in the pulmonary vasculature (8). We show that TGFBI activates NFκB in PEC and enhances NFκB-mediated EC migration via αvβ3 integrins. We further show that TGFBI stimulation increases *Csf3* expression, serving to enhance NO production. Finally, we demonstrate that loss of TGFBI in mice impairs pulmonary vascular development at baseline and severely impairs alveolar and vascular growth in chronic hyperoxia, and that TGFBI expression and localization is aberrant in a preterm lamb model of disrupted alveolarization. In summary, our studies identify a novel axis, whereby developmental expression of TGFBI activates NFκB and promotes pulmonary endothelial angiogenesis during this critical window of vascular development.

Pulmonary angiogenesis is essential for alveolarization, and disrupted angiogenesis contributes to the pathogenesis of bronchopulmonary dysplasia (BPD), the most common complication of premature birth (33). Our lab previously identified the NFκB pathway as an important regulator of pulmonary angiogenesis during alveolarization (8). However, the mechanisms allowing for temporal-specific activation of pro-angiogenic NFκB signaling in the pulmonary vasculature was not known. These results highlight the role of paracrine factors secreted from alveolar myofibroblasts in the creation of a pro-angiogenic niche that activates NFκB to support pulmonary vascular growth during early alveolarization.

By profiling developmental differences in the lung microenvironment, we identified TGFBI as a temporally-regulated protein highly expressed during early alveolarization. TGFBI binds both extracellular matrix (34,35) and integrins (36-39), suggesting a possible role as a bifunctional linker protein that connects cells to the matrix (34). TGFBI mRNA is biphasically altered in the hyperoxia mouse model of BPD (11) and induced during bleomycin-mediated fibrotic lung injury (40). Importantly, single cell RNA sequencing in the developing murine lung identified TGFBI as a highly discriminating gene for myofibroblasts (24). In our study, TGFBI was expressed at the tips of secondary crests, characteristic locations for myofibroblasts (10), concordant with a prior report that identified high expression of TGFBI in the septal tips of a two year-old child, leading the authors to speculate a putative role in alveolar morphogenesis (41).

In other systems, TGFBI is regulated by TGFβ. TGFβ isoforms play a complex role in lung development. Although TGFβ1 is required for lung branching and epithelial differentiation (42), exogenous TGFβ1 inhibits branching of pseudoglandular lung explants (−43). Loss of TGFβ1 does not disrupt lung development (44), but loss of TGFβ3 induces alveolar hypoplasia and extensive intrapulmonary hemorrhage, suggesting role for TGFβ3 in pulmonary vasculature stabilization (45). Impaired alveolarization is also observed in mice with global deletions of Smad3, the downstream effector of TGFβ (46). Taken together these data highlight the importance of precise TGFβ signaling in the correct cells at the right time to support lung development. Further studies will be needed to determine if TGFβ is the primary regulator of TGFBI in the early alveolar lung, however, as a putative downstream effector of TGFβ, our data highlight a role for TGFBI in coordinating alveolar and vascular growth during alveolarization.

TGFBI promotes cell adhesion, migration and proliferation of diverse cell types by interacting via integrins(37, 47, 48). We specifically investigated αvβ3 integrins given their established role in angiogenesis. The αvβ3 integrin is highly expressed by newly forming blood vessels(27). Activation of αvβ3 promotes endothelial migration(49), and blocking αvβ3 inhibits tumor angiogenesis (−50) and impairs lumen formation and vascular patterning in the embryo(51). Further, αvβ3 activates NFκB to promote EC adhesion, survival, and migration(28, 52). Moreover, TGFBI promotes adhesion and migration of human umbilical EC via αvβ3 (39). Concordant with these studies, we found that TGFBI-stimulated PEC migration was blocked by inhibiting either NFκB or αvβ3. Taken together, our data demonstrate that TGFBI promotes PEC migration via αvβ3 to induce pro-angiogenic NFκB signaling.

We next investigated the downstream mechanisms by which TGFBI promotes PEC migration, using RNA-Seq to identify novel TGFBI-regulated genes. One of the few, shared targets in early alveolar and adult PEC was *Csf3* (encoding GCSF), a known NFκB-regulated target gene (30). A well-recognized hematopoetic growth factor (53), GCSF also promotes endothelial migration (54). GCSF is produced by EC stimulated with interleukin-1 (55), an NFκB activator (56), and increases the expression and activation of endothelial nitric oxide synthase (eNOS) to augment NO production (31, 57). NO is produced locally at lamellipodia of migrating human EC, and lung EC from eNOS null mice migrate more slowly and display impaired capillary formation (58-60). In our study, TGFBI increased NO production in PEC, and silencing *Csf3* blocked both TGFBI-mediated NO production and PEC migration. In preterm lambs, prolonged IMV reduced eNOS protein and pulmonary capillary and microvessel abundance (61-63). Taken together, these studies identify *Csf3* as a central downstream mechanism for the pro-angiogenic effects of TGFBI.

Finally, as proof of concept for the importance of TGFBI *in vivo*, we performed studies using TGFBI-/- mice, and a preterm lamb model of impaired alveolarization (11, 18). We showed that TGFBI-/- mice have reduced pulmonary vascular density at baseline, and that chronic hyperoxia markedly exaggerated this vascular phenotype. Further, we found that TGFBI expression was reduced in preterm lambs treated with NRS, consistent with delayed alveolarization observed in this group. Importantly, in preterm lambs maintained with the more injurious, IMV strategy, TGFBI expression was abnormally increased along the thickened septal tips, similar to the abnormal accumulation of elastin and mesenchymal cell proliferation reported (16, 64). These studies suggest that both the correct amount and the correct location of TGFBI is required to optimally support vascular growth. Further, these preclinical studies support recent clinical studies that offer additional evidence of the importance of TGFBI in the developing human lung. In a study of 50 twin pairs affected and unaffected with BPD, rare variants in *TGFBI* were associated with an increased risk for BPD (65). In a subsequent, larger study that employed whole exome sequencing in infants with extreme phenotypes of BPD, rare variants in *TGFBI* were again identified in affected but not unaffected subjects (66). Taken together, these studies provide compelling data to highlight the importance of TGFBI in promoting distal lung development and implicate a role for disrupted TGFBI signaling in the pathogenesis of BPD.

In summary, our data identify a paracrine mechanism by which myofibroblast expression of TGFBI promotes pulmonary angiogenesis through an αvβ3/NFκB axis that increases CSF3-mediated NO production. Given the ability of TGFBI to bind extracellular matrix components highly expressed in the developing lung, local secretion of TGFBI by myofibroblasts may serve to create an angiogenic niche that promotes pulmonary vascular growth along the developing septa. Taken together, our studies identify a novel pathway allowing for myofibroblast-endothelial cross talk, and we speculate that TGFBI dysregulation may contribute to the aberrant pulmonary angiogenesis observe in the setting of impaired alveolarization.

## Supporting information

TGFBI_supplemental Figures

## Acknowledgements

None.

