## Supplementary material for "Developmental Expression of Transforming Growth Factor Induced Protein Promotes NF-Kappa-B Mediated Angiogenesis During Postnatal Lung Development": TGFBI_supplemental Figures

ONLINE DATA SUPPLEMENT

**Methods:**

*Mice.*

Neonatal mice at the early alveolar (postnatal day 6-P6) and adult male and female mice (10-12 weeks) C57BL/6 mice were purchased from Charles River Lab for the isolation of primary pulmonary endothelial cells. TGFBI null mice on a C57BL/6J background have been described previously (E1). Mice were interbred to generate TGFBI homozygotes and WT controls. Genotyping were done using primers 5'-TGTCCGTGCCTAAGTGTGAG-3' and 5'-CAGCAGCAGACCATTTTCAA-3' to detect null allele ~399bp; 5'-TCAACAGCCACAGTGAAAGG-3' and 5'-GCCTGTAACCATTTGCCACT-3' to detect wildtype allele ~520bp. Mice were maintained at 22±2°C and fed a standard diet of rodent chow and water ad libitum.

*Preparation and Analysis of Lung Conditioned Media*

Lung tissue was obtained from C5BL/6 mice at the early alveolar (P6), late alveolar (P16) and adult (8-10 weeks) stages of lung development. The pulmonary circulation was perfused with PBS and the lungs inflated with serum free media. The lung tissue was aseptically removed, minced, and weigh normalized by resuspension at a 4:1 media to tissue (vol/wt) ratio in serum free ECM medium with antibiotics. After 24h, the lung conditioned media (LCM) was collected, cleared by centrifugation, aliquoted and frozen at -80° C until use. To account for mouse-to-mouse variability, LCM from multiple mice were pooled before use in the following experiment.

Protein extracts from all three LCM (P7, P10, Adult) were labeled with separate fluorescent proteins and co-separated on a single gel via IEM 2D-PAGE (Hayward, CA). Fluorescent signals from each pair of two samples were then visualized for a total of three comparisons. The gel images were processed using DeCyder establish the ratios of relative expression between each comparison of samples. Selected spots were subjected to in-gel digestion, and identification via mass spectroscopy. Data were processed using Mascot 2.1 (*http://mascot.bio.nrc.ca/*) or Micromass ProteinLynx Global Server 2.0. Peptide data were searched against the latest SwissProt database for protein identification.

*Isolation of Primary Pulmonary Endothelial Cells (PEC)*

PEC were isolated from P6 or adult C57BL/6 mice, from P6 TGFBI KO or WT mice, as described previously in our lab (E2, E3). Briefly, PECs were obtained from 10-12 mouse lungs by digestion of whole lung tissue with collagenase IA (0.5 mg/ml, Sigma, St. Louis, MO) for 30 min at 37°C, followed by incubating cell lysate for 15 min at RT with anti-CD31-coated magnetic beads (Dynabeads, Invitrogen, Carlsbad, CA), resulting in between 2-3 million PEC after bead selection. PEC isolated by this method were previously characterized by flow cytometry and found to be exclusively CD45- , greater than 95% positive for the endothelial specific marker, CD102 (E2), and greater than 80% demonstrating binding to Griffonia simplicifolia, indicating primarily microvascular EC. PEC were also isolated from mice containing an endothelial cell specific deletion of the NFκB activator IKKβ by crossing IKKβ$^{fl/fl}$ mice (E4) with *Pdgfb*-iCre mice (E5). PEC were then cultured in endothelial growth media (EGM) containing 5% FBS with growth factors (EBM-2; Lonza, Basel, Switzerland) at 37°C in 5% $CO_2$. PEC from passage 0-2 were used for all experiments. In some studies, PEC were treated for 4h with either vehicle (DMSO) or the pharmacologic NFκB inhibitor, BAY11-0782 (2.5 μM), followed by TGFBI (10 μg/ml) treatment for 24h for wound healing assay.

*TGFBI Neutralization and Stimulation*

Neutralization of TGFBI was performed by adding either isotype control antibodies (rat IgG, Cell Signaling Danvers, MA) or anti-TGFBI antibodies (LifeSpan BioScience, Seattle, WA) at a concentration of 4μg/ml prior to diluting the LMC 1:1 in starvation media (EGM + 0.2% FBS). For TGFBI stimulation experiments, cells were stimulated with vehicle (PBS) or recombinant TGFBI (R&D Systems, Minneapolis, MN) at a concentration of 10 μg/ml for all studies.

*Inhibition of Integrin αvβ3 Signaling*

Neonatal or adult PEC (6.5-7 $x10^4$ cells /well ) from passage P1 were starved for 2 h, then anti-αvβ3 integrin antibody (4 μg/ml; Abbiotec, Escondido, CA) or isotype control (Cell Signaling, Danvers, MA) were added to each plate for 2h, followed by stimulation with TGFBI (10 μg/ml) for 24h for wound healing assays. For immunostaining of NFκB/p65, cells were plated and attached on the $22mm^2$ coverslips placed in the 6-well plate overnight, and then stimulated with TGFBI (10 μg/ml) for 30 min before images were taken.

*Immunocytochemistry to Assess NFκB Activity*

PEC were grown on $22mm^2$ coverslips placed in a 6-well cell culture plate until form a thin monolayer. Cells were fixed with 100% ethanol and air-dried. Cells were rehydrated and immunocytochemistry performed as previously described using primary antibodies against p65 (1:200; Santa Cruz Biotechnology, Santa Cruz, CA) overnight at 4° (E2). Fluorescent images were captured and quantified using either a Leica DM5500B Upright or Keyence BZ-X700/BZ-X710 Microscope, and Metamorph Image analysis software (Molecular Devices, Sunnyvale, CA).

*Chemotactic Migration Assays*

Boyden chamber assays were done according to a standard protocol (E6). Starvation media +/- TGFBI were added into the lower chamber, and $1x10^5$ PEC were added to the insert in the upper chamber. Cells were incubated with TGFBI for 8h, at the experimental endpoints cells remaining in the inner side of the insert were removed and the insert was placed in medium containing 8µM Calcein AM for 45 min, trypsinized, and the fluorescence measured at excitation wavelengths of 485nm and 520 nm.

*Endothelial Wound Healing Assays*

PEC (6-7 $x10^4$) were incubated in EGM on 48-well plates for 48h. A linear wound was created using a sterile pipet tip, and the EGM media was then replaced with experimental media detailed in the figure legends. Phase contrast images were captured at time 0 and 24h post-treatment and the percentage of the wound area covered at 24h calculated for each experimental group. Images were taken from at least four wells of each experimental group and data were analyzed using Image J software (NIH).

*Microfluidic Chemotaxis Assays*

PEC migration was also assessed using microfluidic chemotaxis assays as previously described (E7). Microfluidic devices were fabricated using standard soft lithography and micromolding techniques at the Stanford Microfluidics Foundry clean room. The cell culture chamber was adsorbed with fibronectin overnight (10 µg/ml) and rinsed three times with buffer.

PEC ($1 \times 10^6$ cells/mL) suspended in starvation medium were injected into the cell culture chamber and allowed to adhere for ~3h. The inlet and outlet of the cell culture chamber were plugged, and tubing inserted into the inlets of the reagent channels and connected to syringes on a syringe pump (World Precision Instruments, Sarasota, FL). By supplying early alveolar conditioned medium, starvation media + rTGFBI (10μg/ml) or starvation media + VEGF (100 ng/ml) to the source reagent channel and starvation medium to the sink reagent channel (flow rate = 10 nl/min), a stable linear gradient of conditioning molecules is formed across the cell culture chamber. Migrating cells were tracked using time-lapse video microscopy (Zeiss Axiovert 200 microscope, Carl Zeiss AB, Stockholm, Sweden). Cells were imaged every 30 min for 750 min using a phase contrast 10X objective and AxioVision time-lapse software (Zeiss). For each condition, three independent trials were performed with ~60 migratory cell tracks recorded per trial. Cell-tracking analysis was performed using ImageJ software with MtrackJ plug-in. Directional histograms of migrating cells were generated using Chemotaxis and Migration Tool freeware (Ibidi, Munich, Germany), which generates a smoothed histogram for the number of cells migrating within a specific angular trajectory (angular bins of 10 degrees).

*PEC Proliferation Assays*

PEC proliferation was determined by BrdU incorporation assays. Neonatal and adult PEC ($6 \times 10^3$) were plated into each well of a 96-well plate and synchronized with starvation media (0.2% FBS) for 16h, prior to stimulation with experimental media. The incorporation of BrdU was then measured by ELISA at 24h per the manufacturer's protocol (Roche Diagnostics, Mannheim, Germany).

*Western Immunoblot*

Whole cell protein lysates were extracted from lung tissue or PEC using RIPA buffer. Proteins were then subjected to SDS-PAGE, transferred to PVDF membranes, and immunblotting with primary antibodies performed as previously described (E3). The appropriate horseradish peroxidase-conjugated secondary antibody were used to detect the immune-complexes as enhanced chemiluminescence signals on Kodak X-ray films (GE Life Sciences, Piscataway, NJ). Western blots were processed and analyzed by Image J software (NIH).

*Immunofluorescent Staining of Lung Tissue*

Immunostaining was performed on formalin-fixed or frozen murine lung sections using techniques previously described (E2), probed with primary antibodies against CD31 (1:50 for frozen tissue; Abcam, Cambridge, MA; or 1:200 for fixed tissue; Dianova, Germany). TGFBI (sodium azide -free ,1:200; R&D system, Minneapolis, MN), NFκB p65 (1:100 Millipore, Billerica, MA) or von Willebrand factor (1:100; Millipore, Billerica, MA). In similar studies, premature lamb lung tissue was probed with a primary antibody against TGFBI (1μg/μl, Abcam, Cambridge, MA) and anti alpha-smooth muscle actin (1:200, Sigma, St. Louis, MO). After staining, mouse slides were examined using a Leica DM5500 upright and a Micropublisher 5MPixel, color digital camera, using HC Plan Apo 25-mm objectives at 10X and 20X magnification (Leica Microsystems, Buffalo Grove, IL). The number of vWF positive vessels (diameter <100 μm) per high-powered field (HPF) was manually counted on 20X images in a blinded fashion. At least 10 non-overlapping fields per mouse were counted, and the number of vWF positive vessels per HPF averaged to give a single value per mouse. Lamb slides were examined using an inverted Zeiss LSM 780 multiphoton laser scanning confocal microscope (Carl

Zeiss, Germany) with a Zen Black software. Image sizes were set to 1024 × 1024 pixels, and images were taken under the 63X high numerical-aperture oil immersion objective lenses. The excitation wavelength was 488 nm for alpha-SMA, 561 for TGFBI and 405 nm for DAPI.

*Electrophoretic Mobility Shift Assays (EMSA)*

Nuclear proteins were extracted from vehicle and TGFBI-treated early alveolar PEC using the NE-PER kit (Pierce, Rockford, IL). A total of 5ug of protein was used for binding reactions with $\gamma$−32P-labeled oligonucleotides containing the κB consensus sequence (Promega, Madison, WI) in a binding buffer containing 500 ng of salmon sperm DNA, 0.01 U of poly(dI-dC), and 0.5 mM DTT as described previously (E2).

*RNA interference:*

Early alveolar PEC at P6 were transfected with 25 nM of NTC, integrin αV, integrin β3, or Csf3 On-Target Plus SMART pool siRNA (Dharmacon, Lafayette, CO; Thermo Fisher Scientific, Lafayette, CO) using Lipofectamine 2000 (Invitrogen) for 6h as previously described (E8). The groups of PEC were then recovered for 42h, prior to use for Western immunoblot. In separate studies, siRNA-transfected PEC were stimulated with vehicle or recombinant TGFBI (10 μg/ml, R&D system, Minneapolis, MN) for 24h then followed by endothelial wound healing assays to assess PEC migration as described above.

*RNA-Seq Analysis*

Early alveolar and adult PEC were cultured in EGM medium until passage 2, then starved overnight, prior to vehicle (PBS) or rTGFBI stimulation for 4h. Total RNA from vehicle and rTGFBI-treated cells were extracted using RNeasy Mini kit (Qiagen, Germantown, MD), and RNA-sequencing performed by Quick Biology (Pasadena, CA). Briefly, RNA integrity was confirmed by Agilent Bioanalyzer 2100, and libraries were prepared according to KAPA Stranded mRNA-Seq poly(A) selected kit (KAPA Biosystems, Wilmington, MA). Final library quality and quantity was analyzed and 150 bp paired end reads sequenced on Illumina HighSeq 4000 (Illumina Inc., San Diego, CA). Reads were mapped to the latest UCSC transcript set using Bowtie2 version 2.1.0 (E9) and the gene expression was estimated using RSEM v1.2.15 (E10). TMM (trimmed mean of M-values) was used to normalize the gene expression. Differentially expressed genes were identified using edgeR (E11). Genes showing altered expression with $p < 0.05$ and more than 1.5-fold change were considered differentially expressed.

*Quantitative real time PCR*

Total RNA was isolated from PEC treated with vehicle or rTGFBI using the RNeasy Mini kit (Qiagen, Germantown, MD). RNA (2 μg) was reverse-transcribed using Superscript III (Invitrogen), and qPCR was performed using TaqMan primers (Applied Biosystems, Carlsbad, CA) as described previously (E12). Quantification of target gene expression was performed using the Delta–Delta CT method and normalized to the expression of 18s.

*Measurement of Nitric Oxide (NO) Production in PEC.*

PEC were grown to 80% confluence, starved for 2h, and then stimulated with vehicle or rTGFBI for 4 or 24h. NO production was determined by loading the cells with 4-amino-5-

methylamino-2',7'-difluorofluorescein diacetate (DAF-FM) (10 μM) (Invitrogen Molecular Probes) (E13) for 40 min at 37 °C, prior to immunofluorescent imaging as previously described (E14). Images were captured using a Keyence BZ-X700/BZ-X710 Fluorescence microscope with a cooled CCD camera (Keyence Corporation of America, IL). Fluorescence intensity of the images was quantified using ImageJ software (NIH). The ratio of total DAF fluorescence in the image to the total nuclei was used to normalize the variation in cell numbers in a particular field. Each experiment was repeated at least 3 times with more than six images taken per group for quantification.

*Murine Model of BPD Induced by Chronic Hyperoxia*

Litters of newborn $TGFBI^{+/+}$ and $TGFBI^{-/-}$ pups at P0 were split into two groups and maintained in either room air (normoxia) or 80% $O_2$ (hyperoxia) in a BioSpherix chamber (BioSpherix, Parish, NY) for 14 days, an established experimental model that recapitulates the impaired alveolarization observed in bronchopulmonary dysplasia (E15). Dams were rotated every 24h to prevent oxygen toxicity. Pups were euthanized at P14 and lungs fixed with 4% paraformaldehyde at 25 cm $H_2O$ pressure for histology. Morphometric analysis (radial alveolar count and airspace area and size) was performed as previously described (E2, E3).

*Preterm Lamb Model of Impaired Alveolarization*

The methods for delivery and continuous management of chronically ventilating preterm lambs are reported (E16-E19). Briefly, time-pregnant ewes that carried one fetus or twin fetuses at 132 ± 2 d of gestation (term ~150 d gestation) were used. The pregnant ewes were given dexamethasone phosphate ~36 h before operative delivery. At delivery, all preterm fetal lambs were intubated, administered Survanta before operative delivery, resuscitated by invasive mechanical ventilation (IMV), and given daily with postnatal caffeine citrate to stimulate respiratory drive. At ~3 h of age, the preterm lambs were randomized to continue IMV or transitioned to noninvasive respiratory support (NRS), provided by high-frequency nasal ventilation as previously described (E18) for a total of 21d. Control lambs were born at term and lived in the laboratory at the University of Utah Health Sciences Center.

Control and preterm NRS lambs were intubated and reconnected to the ventilator to maintain lung inflation when the chest was opened to remove the lungs. The whole left lung was insufflated with 10% buffered neutral formalin (static pressure of 25 $cmH_2O$), and immunofluorescence of paraffin-embedded lung sections performed as described above.

*Statistics*

All data are presented as mean ± SEM. Statistical differences between two groups were determined by Student's t-test. Comparison between more than two groups was analyzed by One-Way (for one independent variable) or Two-Way ANOVA (for two independent variables), followed by Bonferroni Multiple Comparison post-hoc analysis. A P value of $\leq 0.05$ was considered statistically significant.

*Study Approval*

Protocols for the murine and lamb studies adhered to American Physiological Society/US National Institutes of Health guidelines for humane use of animals for research and were prospectively approved by the Institutional Animal Care and Use Committee at Stanford University and the University of Utah Health Sciences Center.

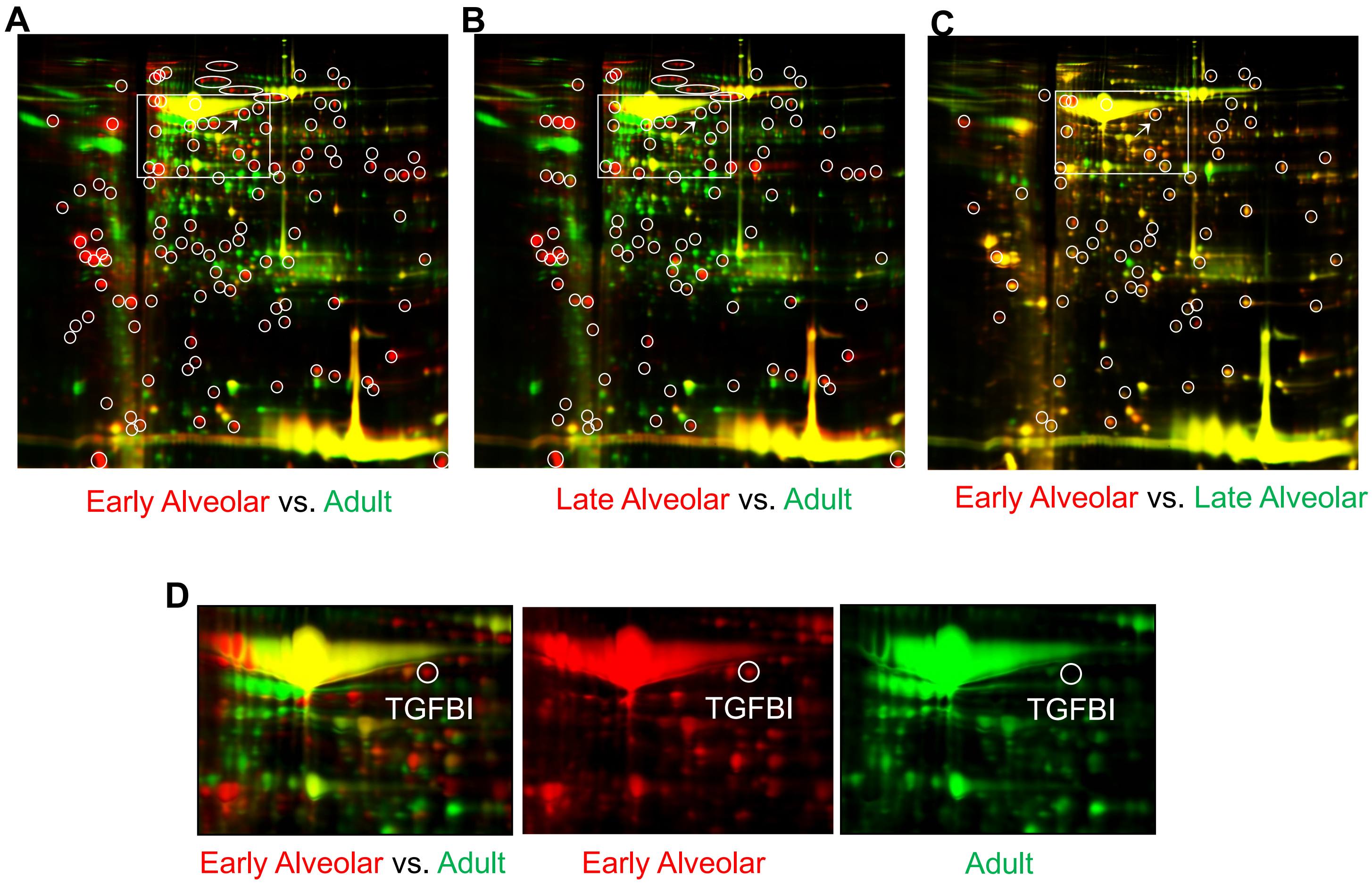


**Figure E1: Proteomic analysis of lung conditioned media identifies TGFBI as a factor uniquely secreted by the early alveolar lung.** Three separate fluorescent images of the same 2D gel after electrophoresis of proteins contained within the three LCM, imaged to compare the relative expression in each of the three comparison pairs: (A) early alveolar vs. adult; (B) late alveolar vs. adult; (C) early alveolar vs. late alveolar. In each comparison, proteins of one media are visualized in red and the protein of the other media visualized in green. Circled proteins were present in the early or late alveolar LCM but absent in the adult (Fig. 2A and B), and also increased in the early alveolar vs. the late alveolar LCM (Fig. 2C). Arrow points to the spot corresponding to TGFBI. (D) Enlarged image of the section of gel, containing the protein spot corresponding to TGFBI.

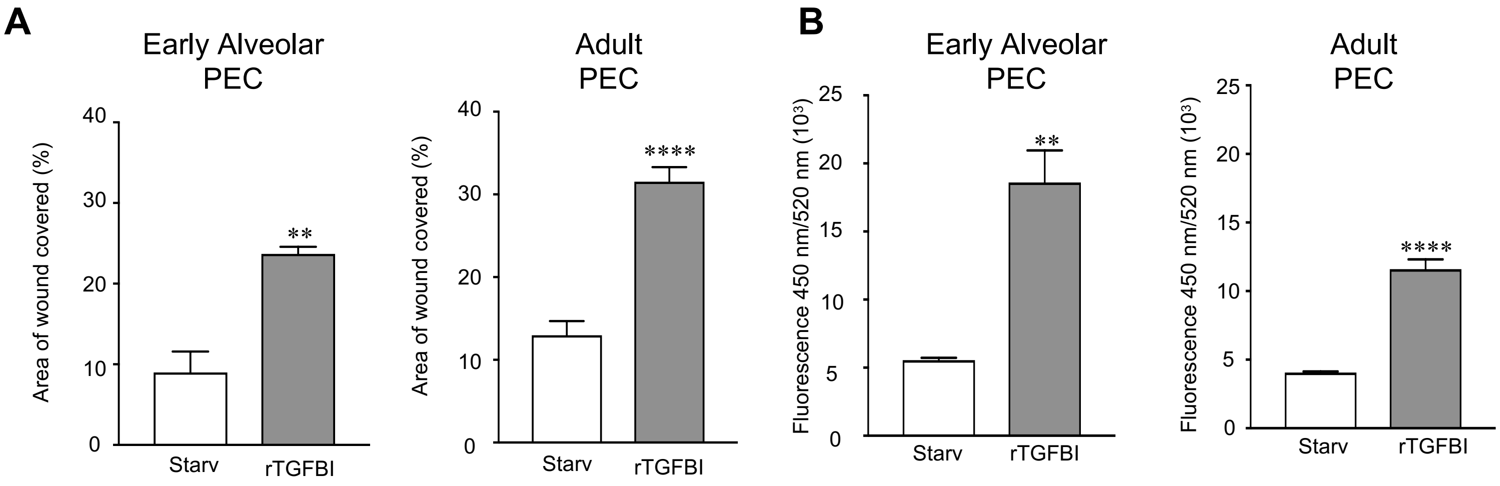


**Figure E2: TGFBI is sufficient to promote early alveolar and adult PEC migration.** (A) Endothelial scratch assays were performed using early alveolar and adult PEC incubated with starvation media or starvation media containing rTGFBI and the percent scratch area covered at 24h calculated. **P<0.01 and ****P<0.0001 vs. starvation with n=3 per group. (B) Boyden chamber assays in early alveolar and adult PEC incubated with starvation media or starvation media containing rTGFBI for 5h. **P<0.01 and ****P<0.0001 vs. starvation media with n=3 per group.
